# A *cis*-regulatory sequence of the wing selector gene, *vestigial*, drives the evolution of scaling relationships in *Drosophila* species

**DOI:** 10.1101/2022.06.23.497339

**Authors:** Keity J. Farfán-Pira, Teresa I. Martínez-Cuevas, Timothy A. Evans, Marcos Nahmad

## Abstract

Scaling between specific organs and overall body size has long fascinated biologists because they are a primary mechanism through which organismal shapes evolve. Yet, the genetic mechanisms that underlie the evolution of allometries remain elusive. Here we measured wings and tibia lengths in four *Drosophila* species (*D. melanogaster, D. simulans, D. ananassae*, and *D. virilis*) and show that the first three of them follow a single evolutionary allometry. However, *D. virilis* exhibits a divergent wing-to-tibia allometry due to a dramatic underscaling of their wings with respect to their bodies compared to the other species. We asked whether the evolution of this scaling relationship could be explained by changes in a specific *cis*-regulatory regulatory region of the wing selector gene, *vestigial* (*vg*), whose function is broadly conserved in insects and its expression pattern determines wing size in *D. melanogaster*. To test this hypothesis directly, we used CRISPR/Cas9 to replace the DNA sequence of the predicted Quadrant Enhancer (*vg*^*QE*^) from *D. virilis* for the corresponding *vg*^*QE*^ sequence in the genome of *D. melanogaster*. Strikingly, we discovered that *D. melanogaster* flies carrying the *D. virilis vg*^*QE*^ sequence have wings that are much smaller with respect to controls, partially rescuing the wing-to-tibia ratio observed in *D. virilis*. Our results show that this *cis*-regulatory element in *D. virilis* contributes to the underscaling of wings in this species. This provides evidence that scaling relationships may be unconstrained and may evolve gradually through genetic variations in *cis*-regulatory elements.

**Summary statement:** Using CRISPR/Cas9 replacement of a *cis*-regulatory element, this study suggests that changes within the *vestigial* Quadrant Enhancer sequence are responsible for the evolution of wing allometries in *Drosophila* species.

## INTRODUCTION

Genetic and environmental factors contribute to morphological diversity across related species (Carroll, 2000). For instance, evolution of morphogenetic traits may be driven by environmental cues followed by selection of specific genetic variations in a population (Uller et al., 2018). The search for genetic variations that account for phenotypic changes across species has attracted the attention of evolutionary biologists for decades, but only recently, genome editing technologies have enabled us to test some predictions directly (Ryan and Farley, 2020). One way to study the genetic contributions to related-species diversity is to investigate how changes in the genome could explain the evolution of certain organs to whole-body scaling relationships (Evans, 2017; Van Belleghem et al., 2020).

To investigate scaling relationships, we will use the concept of allometry, this is, the evaluation of the relationship between specific traits. Allometries have been used to analyze morphological changes that give rise to diversity and phenotypic variation in ecology and evolution (Esquerré et al., 2017; Galicia-Mendoza et al., 2021; Gomes Rodrigues et al., 2018; Shingleton, 2010). Depending on how scaling relationships are studied, allometries can be classified as *ontogenic* (scaling relationships during development and growth within an organism (Esquerré et al., 2017; Simons and Frost, 2021)); *static* (scaling relationships across a population of organisms in a specific stage of development (Shingleton et al., 2008)), and *evolutionary* (scaling relationships across individuals of different species (Tidière et al., 2017)). Mathematically, the scaling of any two traits or characters *x* and *y* within an organism can be modeled using the growth law:

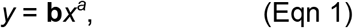

where **a** (known as the allometry coefficient) and **b** are parameters that are fitted to data measurements of (*x, y*) pairs. Note that Eqn 1 is approximately linear only when **a**≈1, and therefore, it is usually easier to employ a logarithmic transformation to get a linear model, regardless of the value of **a**:

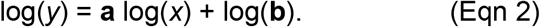

In Eqn 2, the transformed variables log(*x*) and log(*y*) are fitted to a linear transformation using standard linear-fitting procedures, where **a** is the slope of the line and log(**b**) is the intercept (Gayon, 2000; Shingleton 2010). Using this formalism, allometries have been used to understand how environmental conditions (such as temperature, nutrition and rearing density) influence natural variation and trigger variability in scaling patterns (Shingleton et al., 2009; Weber, 1990), or investigate the relationships underlying morphological evolution across different species (Pélabon et al., 2014; Shingleton, 2010). However, the genetic mechanisms underlying the evolution of allometric relationships in a group of species are little understood.

One way to understand the genetic basis of phenotypic evolution is to focus on key candidate genes involved in the developmental process of interest (True and Haag, 2001). Based on this candidate gene paradigm, allometries may evolve acting on inputs of the gene regulatory networks that control the expression pattern of a key selector gene (Fig. 1A). Alternatively, evolution could act on genetic sequences that alter the properties of the protein that is produced by the selector gene (e.g., stability, degradation, or its ability to bind DNA or other proteins; Fig. 1B). Finally, evolution may be driven by changes in *cis*-regulatory elements (promoters, enhancer, or silencers) of the selector gene (Rebeiz and Tsiantis, 2017; Ryan and Farley, 2020) (Fig. 1C).

**Figure 1.**
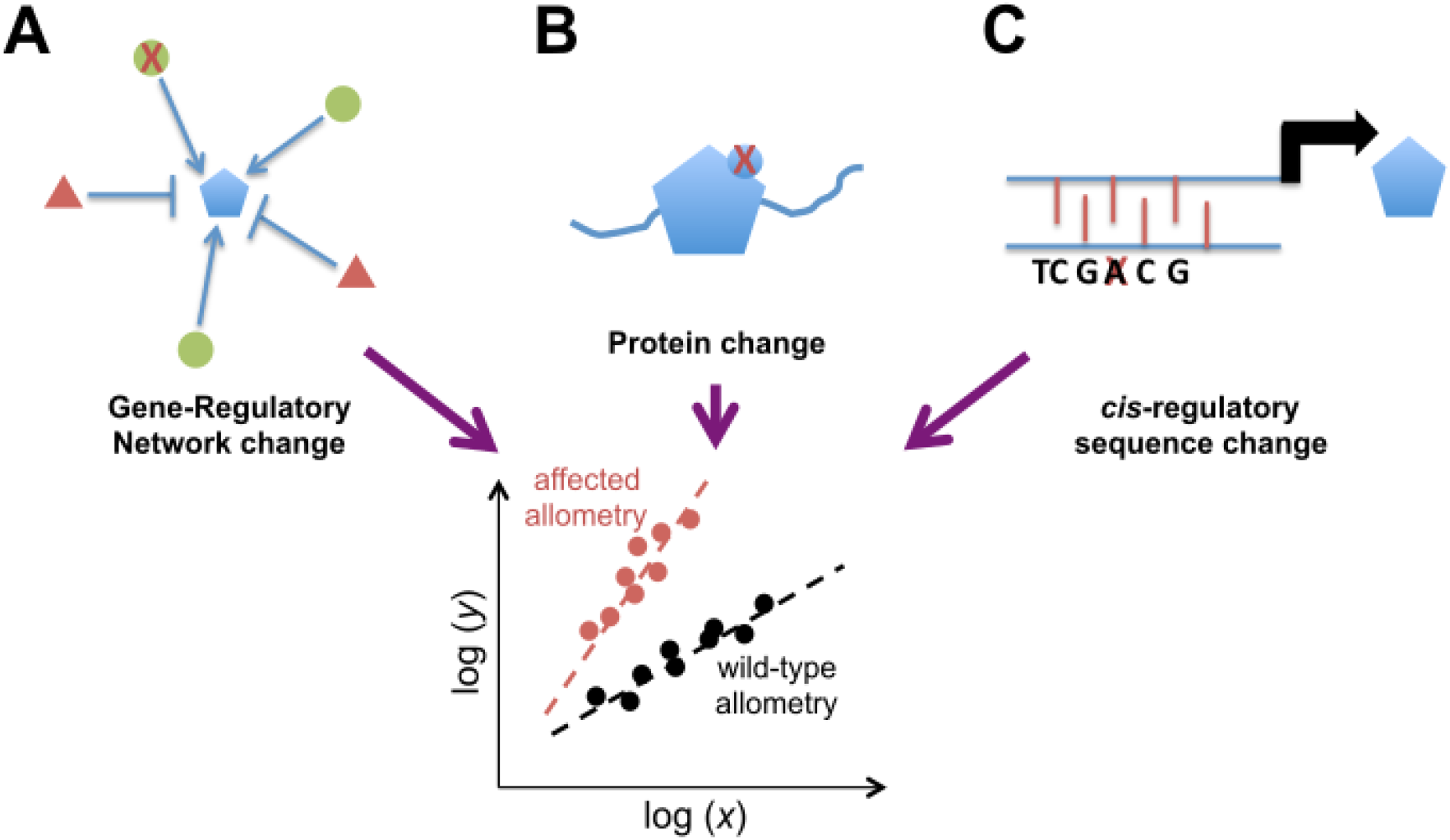
Selector-gene mechanisms for allometric evolution. Scaling relationships between two traits *x* and *y* may evolve by affecting (red ‘X’) a selector gene (represented by a blue pentagon) in three different ways: (A) By altering the expression or effect of an activator (green circle) or a repressor gene (red triangle). (B) By affecting the stability, function, or post-translational modification of the selector-gene protein product. (C) Causing a modification in the *cis*-regulatory region of the selector gene.

In holometabolous insects (Diptera, Lepidoptera, Coleoptera and Hymenoptera), *vestigial (vg)* is an evolutionarily conserved wing selector gene (Abouheif and Wray, 2002; Clark-Hachtel et al., 2013; Nel et al., 2013). In *Drosophila melanogaster*, loss of *vg* function leads to the development of an adult fly with only vestiges of wing tissue (Williams et al., 1991; Williams et al., 1993), while Vg overexpression induces wing-like transformations (Baena-López and García-Bellido, 2003; Halder et al., 1998; Kim et al., 1996; Williams et al., 1994). Vg is part of a transcription factor complex that has an important role in larval wing and haltere development (Simmonds et al., 1998). In this complex, Vg interacts with the transcription factor Scalloped (Sd) to activate the transcription of wing-differentiation genes (Delanoue et al., 2004; Halder et al., 1998; Williams et al., 1991). *vg* expression is controlled through the influence of at least two regulatory regions: the Boundary Enhancer (*vg*^*BE*^) which drives expression along the Dorsal-Ventral border of the wing disc that will become the adult wing margin, and the Quadrant Enhancer (*vg*^*QE*^), that contributes to Vg protein expression in the rest of the wing pouch (Klein and Arias, 1999; Williams et al., 1994; Zecca and Struhl, 2007a).

In order to ask whether allometric relationships evolve through genetic changes in a key selector gene (Fig. 1), here we investigate how wing sizes scale relative to tibia sizes (used as a proxy to whole-animal size) in four *Drosophila* species: *D. melanogaster, D. simulans, D. ananassae*, and *D. virilis*. While there are some deviations in each of the allometric relationships among all species, we immediately noticed that *D. virilis* behaves very different from the other three species. Particularly, the wings of *D. virilis* do not follow the scaling relationship expected by the evolutionary allometry defined by the other species. Thus, we asked whether this scaling relationship evolved through *cis*-regulatory elements within the *vg* gene (Fig. 1C). To address this, we cloned the corresponding sequences of the *vg*^*QE*^ and used CRISPR/Cas9-based replacement of these sequences into the *D. melanogaster* genome. Strikingly, in CRISPR-edited animals, where the *vg*^*QE*^ of *D. melanogaster* was replaced by the corresponding *vg*^*QE*^ of *D. virilis*, wings underscale with respect to what would be expected in wild-type *D. melanogaster* flies. These results suggest that the *vg*^*QE*^ sequence plays an important role in the evolution of wing-to-body scaling relationships in *Drosophila* species.

## MATERIAL AND METHODS

### *Drosophila* strains

The following *Drosophila* strains were used: *D. melanogaster* [Meigen, 1830] Samarkand strain (RRID:BDSC_4270 - Bloomington Drosophila Stock Center); *y,w* strain of *Drosophila melanogaster*, provided by Dr. Fanis Missirlis (Cinvestav, Mexico); *D. simulans* [Sturtevant, 1919] (14021-0251.261 - Drosophila Species Stock Center); *D. ananassae* [Doleschall, 1858] (14024-0371.00 - Drosophila Species Stock Center); *D. virilis* [Sturtevant, 1916] (15010-1051.87 - Drosophila Species Stock Center); w, ms1096-Gal4 (Bloomington Drosophila Stock Center #RRID:BDSC_8860); UAS-*vg*^*RNAi*^ (Vienna Drosophila Resource Center #16896); nub-Gal4 (Bloomington Drosophila Stock Center #RRID:BDSC_38418); *w; Sco/CyO, RFP* (from Timothy A. Evans); *y, w*, nos-Cas9 (Bloomington Drosophila Stock Center #RRID:BDSC_54591), *D. melanogaster (vg*^*QEDmel*^) and *D. melanogaster (vg*^*QEDvir*^; generated in this study). Adult flies were kept at 25°C in vials containing standard *Drosophila* food. All crosses were carried out at 25 °C.

### Sequence alignment and phylogenetic analysis

The CDS sequences analyzed in this study are available in FlyBase and NCBI database (*D. melanogaster* FBgn0003975 and NM_078999.4, *D. simulans* XM_016167959.2, *D. yakuba* XM_002090946.3, *D. erecta* XM_001975804.3, *D. ananassae* XM_001959506.4, *D. virilis* XM_032435676.1, *D. mojavensis* XM_002006424.4, *D. grimshawi* XM_001986428.2) were used in multiple alignments using MEGA X software version 10.2.4 (Kumar et al., 2018). Phylogenetic tree was developed using a Bayesian model in MrBayes software version 3.2.7a (Ronquist and Huelsenbeck, 2003). The analysis was performed with 23,000 generations, sampling phylogenies each 1000 generations and to reach a SD = 0.013933.

Synteny of *vg* gene in *D. melanogaster, D. simulans, D. ananassae* and *D. virilis* was evaluated through information of NCBI database (https://www.ncbi.nlm.nih.gov/) and OrthoDB database (https://www.orthodb.org/). The search was limited to the fourth intron of the gene, and posteriorly paired/local alignments using .fasta format through Smith Waterman algorithm (Smith and Waterman, 1981) in EMBOSS Water (Ubuntu 18.04 https://launchpad.net/ubuntu/bionic/+package/emboss) were developed (*D. melanogaster* QE *vs. Drosophila* species introns): length of each sequence was delimited based on *D. melanogaster* reference sequence, followed by multiple alignments using Clustal Omega from EMBL-EBI (Madeira et al., 2022). Sequence of QE reported for *Drosophila melanogaster* strain Samarkand (GenBank ID: FJ513071.1) was used as a template for searching of QE sequences for *D. simulans, D. ananassae*, and *D. virilis*.

### Allometries

Adults were separated by sex (data in the main figures, Fig. 3 and 5, correspond to females; but equivalent results were obtained for males in Fig. S2 and S4, respectively), using a stereoscopic microscope (Nikon SMZ800) and preserved in 1ml 70% ethanol for dehydration for at least 12 hours for further analysis. Each specimen was dissected in 15μl of ethanol 50% to obtain right and left wings, and right and left front legs, which were mounted in microscope slides using a stereoscopic microscope (Nikon SMZ800).

Wings and legs were photographed in a bright-field binocular microscope (Nikon Eclipse Ci) attached to a camera (ProgRes® CT5, Jenoptik) using software ProgRes® Capture Pro-2.9. Measurements of proximal distal length (see Fig. 3A-B) were performed using ImageJ software (https://imagej.nih.gov/ij/download.html) (Schneider et al., 2012) after the corresponding calibration for 4X objective (Distance in pixels: 100.501; known distance: 0.1; pixel aspect ratio: 1.0, unit of length: mm). Each point in the allometry graphs correspond to the average measurements of two wings and two legs of the same animal. We used ordinary least squares (OLS) to estimate the regression line (after a log10 transformation) and slope for each sex.

### gRNA Design and construction of gRNA plasmids

Target sites were designed using flyCRISPR online tool CRISPR Optimal Target Finder (https://flycrispr.org) (Gratz et al., 2014; Iseli et al., 2007). Cloning was performed into the pCFD4-U6:1_U6:3tandemgRNAs vector (Addgene plasmid # 49411; (http://n2t.net/addgene:49411;RRID:Addgene_49411), that allows tandem expression of two gRNA sequences (Evans, 2017; Port et al., 2014), through PCR amplification using pair of primers 2 and 3 (Table S1) with a 2X Phusion Flash PCR Master Mix (Thermo Scientific, Cat. # F548S). Gibson Assembly (New England Biolabs, Cat. # E2611) was performed with PCR products and pCFD4 *Bbs*I (New England Biolabs, Cat. # R0539L) digested vector. All cloning products were confirmed by DNA sequencing prior to injection.

### Construction of QE donor plasmids

The donor plasmid was assembled using overlap extension PCR (Heckman and Pease, 2007). We first designed a working base vector composed of four PCR fragments, two that were derived from plasmid backbone of pHD-DsRed (Addgene plasmid # 51434; https://www.addgene.org/51434/) (primer pair 835, 836), DsRed fluorescent coding region (primer pair 837, 838), and two derived from genomic DNA of *D. melanogaster*, left homologous arm (LHA2) (primer pair 839, 840) and right homologous arm (RHA1) (primer pair 841, 842). We then assembled the LHA2-DsRed-RHA1 fragments in one round of PCR (primer pair 839, 842) and then combined this product with the plasmid backbone fragment in a second PCR to obtain the whole base vector as a linear fragment: pHD-DsRed-LHA2-DsRed-RHA1 (primer pair 837, 843). The PCR product was transformed directly into competent *E. coli* and circularized *in vivo*. To insert the specific QE sequences, we amplified by PCR the sequences from genomic preparations of *D. melanogaster* (primer pair 844, 845) and *D. virilis* (primer pair 844, 846) and assembled with the base vector through overlap extension PCR. All the components of the donor vector including the QE, were sequenced prior to injection.

### Identification of CRISPR-modified alleles

The *vg*^*QE*^ gRNA plasmid was co-injected with the QE homologous donor plasmids into nos-Cas9 embryos by Rainbow Transgenic Flies (Camarillo, CA, USA). Individuals recovered from injected embryos were crossed to w; *Sco/CyoRFP* flies. CyORFP or Sco flies from this cross (potentially carrying *vg*^*QE*^ HDR alleles) were screened for red fluorescent eyes (indicating genomic integration of DsRed sequences carried on the HDR donor plasmid; see Fig. S3). Fluorescent-eye flies were then crossed back individually to w; *Sco/CyORFP* to generate balanced stocks. Confirmation of the inserted sequences was tested by PCR using primers 20, 23, 41, 44, 47, and 53 (Table S1) (see Figure S3). PCR-confirmed balanced stocks with the modified alleles were also confirmed through genomic DNA sequencing.

### Statistical analysis

Allometry data was analyzed using R version 4.1.2 (https://www.r-project.org/) and RStudio (https://www.rstudio.com/). ANCOVA tests for slope comparisons were carried out using lm() and anova() commands. Wing and tibia lengths, and wing/tibia ratios were analyzed using GraphPad Prism 8.0.1 Software (https://www.graphpad.com/scientific-software/prism/), through one-way ANOVA with a *p-value*<0.05 to define significance.

## RESULTS

### Construction of a phylogenetic tree using the *vg* gene

In order to study wing-to-body allometric relationships in *Drosophila* species, we selected four representative species based on the following criteria. Since *vg* is evolutionarily conserved as a wing selector gene in insects (Abouheif and Wray, 2002; Clark-Hachtel et al., 2013; Clark-Hachtel et al., 2021; Zhang et al., 2021), we first used the *vg* coding sequence (CDS) from eight species closely related to *D. melanogaster* to construct a phylogenetic tree (Fig. 2). We then choose two species within the *melanogaster* subgroup (*D. melanogaster* and *D. simulans*), another species outside of this subgroup but still belonging to the *melanogaster* group (*D. ananassae*), and another species from an outside group (*D. virilis*). Wings in these species are similar in shape and pattern (Kumar et al., 2022), but animals across these species vary considerably in whole-body size (Morin et al., 1996). This suggests that the evolution of wing-to-body allometries may be investigated by comparing these species.

**Figure 2.**
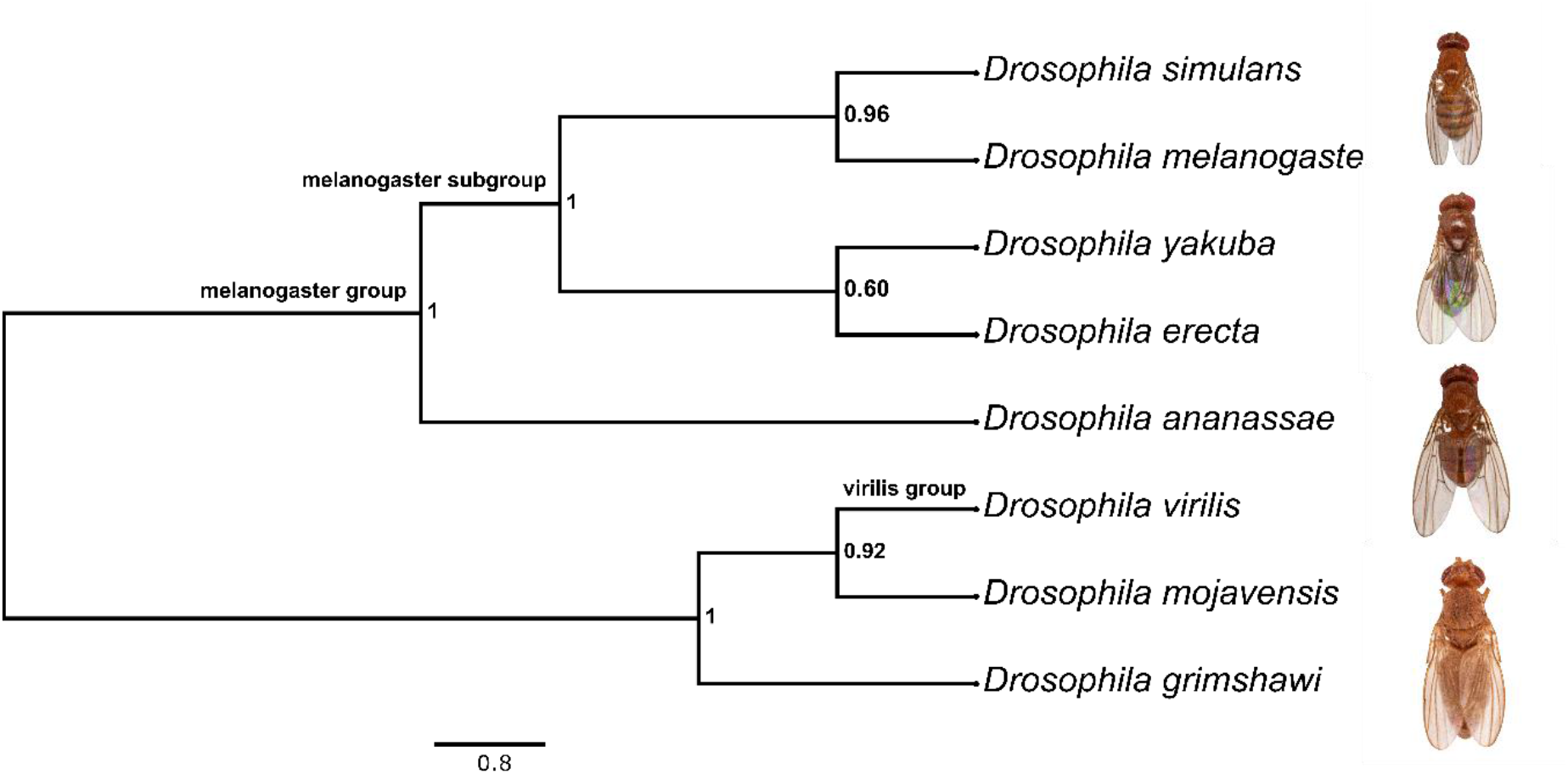
Phylogenetic tree based on the *vg* gene in *Drosophila* species. Bayesian phylogenetic tree using the CDS sequence of the *vg* gene of several *Drosophila* species. Numbers at the nodes correspond to the posterior probability, which supports that the association made by the model is correct. Maximum value of posterior probability is 1. The scale indicates substitution per site. Photos correspond to representative individuals of the species used in this study.

**Figure 3.**
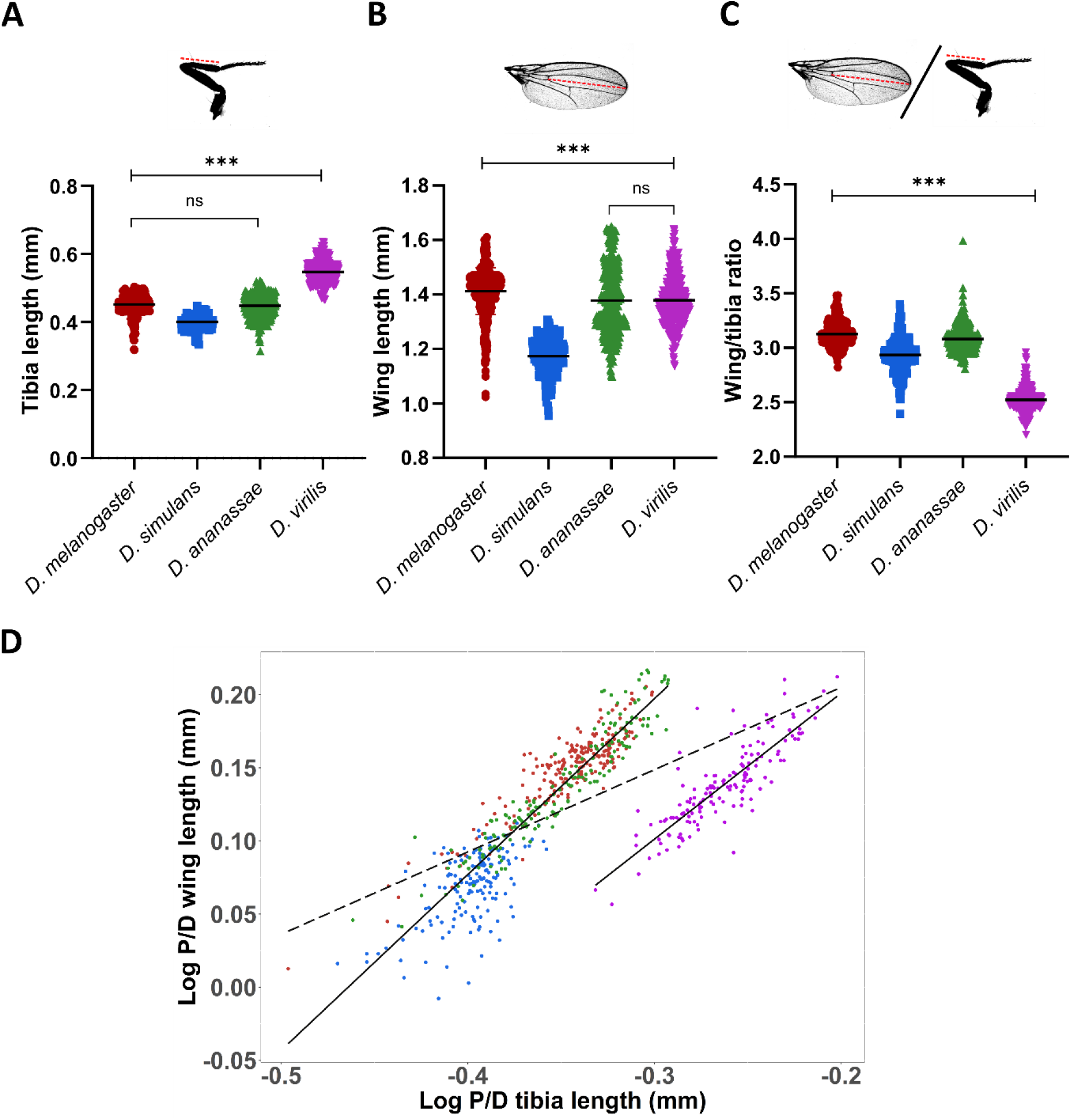
Wing-to-tibia allometric relationships reveal underscaling of wings in *D. virilis*. (A, B) Proximal-distal lengths (red dotted lines) of tibias (from anterior legs, A) and wings (B) were measured and compared in female flies (equivalent results were obtained for males; see Fig. S2) of the four species. Each dot in the distribution is the average of each pair of tibias and wings, respectively. (C) Wing-to-tibia ratios from measurements in each side (left and right) of an animal were computed and statistically compared. (A-C) Mean values are shown. One-way ANOVA analysis using multiple comparisons with Tukey test, \*\*\**p<0*.*001, ns =* Not Statistically Significant (*p>0*.*05*). (D) Wing-to-tibia evolutionary allometric relationships. Black line corresponds to the best-fit linear allometric relationship (equation 2) considering *D. melanogaster, D. simulans*, and *D. ananassae* in a single group (R^2^=0.856). Black-dotted line shows the allometric relationship if all four species were considered (R^2^=0.4738). Experiment replicated two times in the laboratory. *D. melanogaster n*= 224, *D. simulans n*=157, *D. ananassae n*=169, *D. virilis n*=156.

### Wings underscale in *D. virilis* relative to the other *Drosophila* species

Although several measures of whole-body size (e.g., weight) have been used in *Drosophila* studies (Mirth and Shingleton, 2012; Morin et al., 1996; Stillwell et al., 2011), they are sometimes difficult to measure individually or are not directly comparable to wing measures (because they are measured in different units, e.g. weight *vs*. wing area). Therefore, we looked for a trait that could serve as a reference to draw allometric relationships with the wings. The length of the tibia in anterior legs (Fig. 3A top) scale with overall body size in several Dipterans (Krause et al., 2019; Reigada and Godoy, 2005) and can be easily measured and directly compared to the proximal-distal (PD) length of the adult wing (Fig. 3B top; Material and Methods). In addition, legs are developmentally independent to wings since they, unlike wings, are not affected by changes in *vg* expression (Fig. S1A, B). Relative to *D. melanogaster*, mean values of tibia and wing lengths follow the same trend in *D. simulans* and *D. ananassae*; for instance, smaller tibias in *D. simulans* also correspond to smaller wings (Fig. 3A, B and Fig. S2A, B). But *D. virilis* individuals, which are much bigger animals and exhibit a correspondingly larger tibia length (relative to *D. melanogaster*) do not scale their wings accordingly (magenta distributions in Fig. 3A, B and Fig. S2A, B). This mean-value underscaling of wings in *D. virilis* is also revealed by comparing the wing-to-tibia ratio (Fig. 3C and Fig. S2C). At the population level, animals of each species display nearly isometric scaling, *i*.*e*., individual allometries display slope values close to 1 (Fig. S2E). However, while a single evolutionary allometry fit the data of *D. melanogaster, D. simulans*, and *D. ananassae* altogether (solid line through these data points in Fig. 3D and Fig. S2D; Table 1 and Table S2), the data of *D. virilis* do not fit in this scaling relationship (dotted line in Fig. 3D and Fig. S2D; Table 1 and Table S2). This conclusively shows that *D. virilis* underscale the size of its wings, relative to the other species.

**Table 1.**
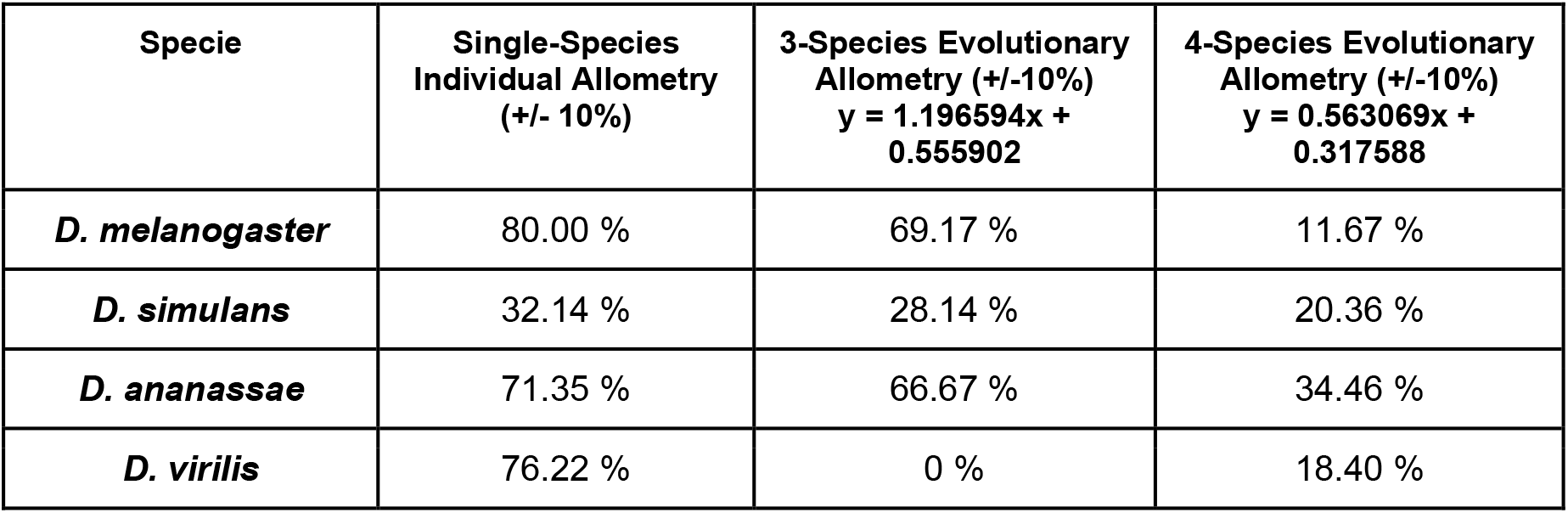
Percentage of single-species data that are explained by the allometry model.

### Bioinformatic identification of regulatory elements within the *vg*^*QE*^ sequence

We considered the possibility that wing-to-tibia scaling was controlled by *cis*-regulatory elements of the *vg* gene (Fig. 1C). From the two intronic *cis*-regulatory elements that drive *vg* expression (Fig. 4), we focused on the *vg*^*QE*^ because it controls expression in most of the wing pouch (compare cyan and magenta patterns in Fig. 4A), and previous work has implicated this element in wing growth (Muñoz-Nava et al., 2020; Zecca and Struhl, 2007a). We first examined the sequence of the reported *vg*^*QE*^ (Dworkin et al., 2009) GenBank ID: FJ513071.1) in *D. melanogaster* and noticed that there is a new Scalloped-interaction-domain (SID) upstream of the reported QE sequence (Fig. 4B; Farfán-Pira et al., 2022). In order to find whether a *cis*-regulatory element also exist in the other three *Drosophila* species, we aligned the sequences of the whole fourth intron of each four species selected in Fig. 2. The alignment recovers sequence conservation, phylogenetic analysis, and perfect-match SIDs (arrowhead) in all four species. In addition, we found an extra SID in *D. ananassae* (Fig. 4C,D). This analysis allowed us to predict a *vg*^*QE*^ in each of these species.

**Figure 4.**
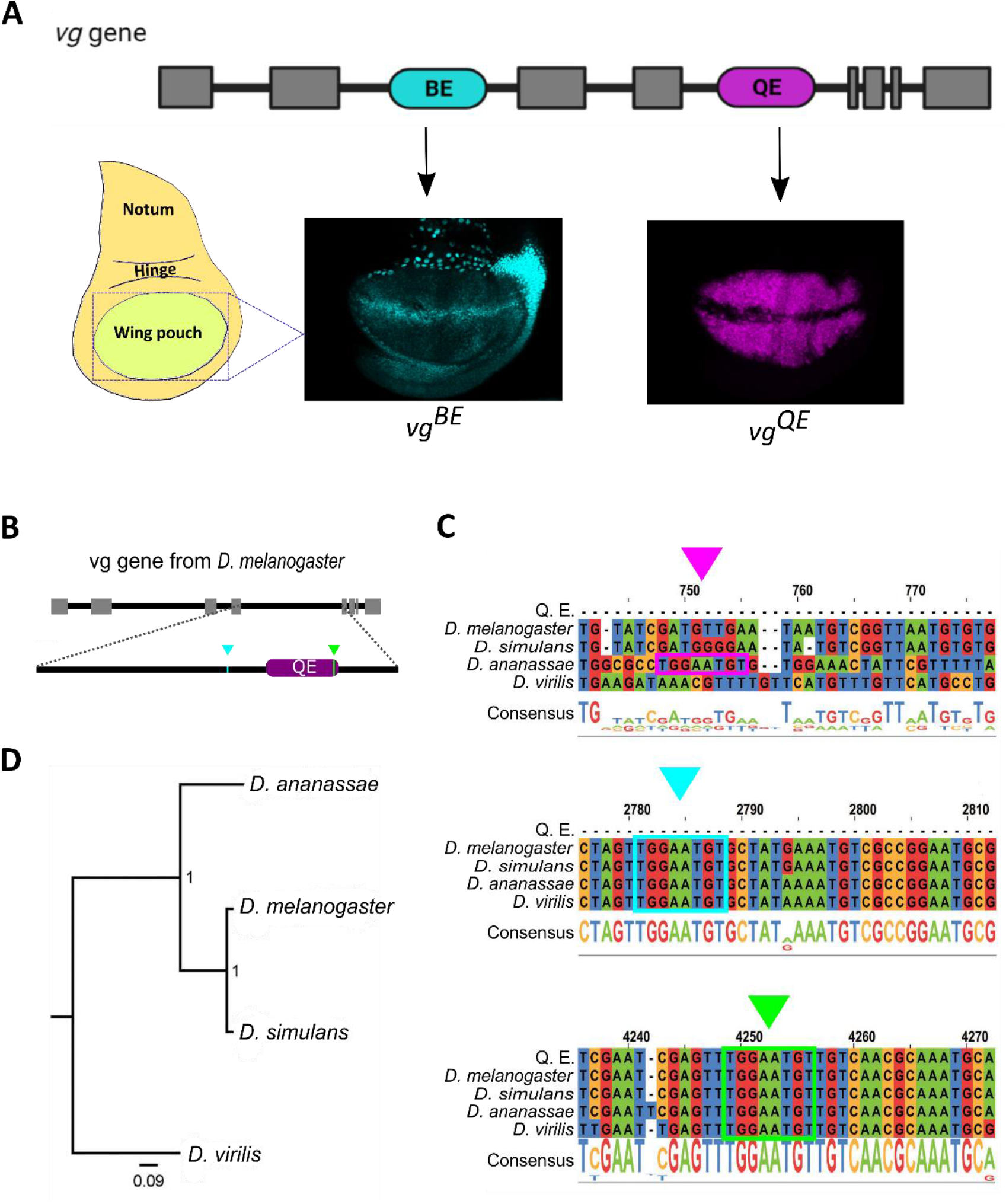
Regulatory sequences of the vg gene from *Drosophila*. (A) Expression pattern of the *cis*-regulatory sequences on the vg gene of the wing disc from *D. melanogaster* (Taken from Farfán-Pira et al. 2022). (B) Schematic representation of Scalloped Interaction Domain (SID) sites (arrowheads) of vg gene from *D. melanogaster*. (C) Identification of SID sites on the 4th intron of vg gene from *D. melanogaster, D. simulans, D. ananassae* and *D. virilis*. (D) Bayesian phylogenetic tree using the sequences of the 4th intron of the *vg* gene from *D. melanogaster, D. simulans, D. ananassae* and *D. virilis*.

**Figure 5.**
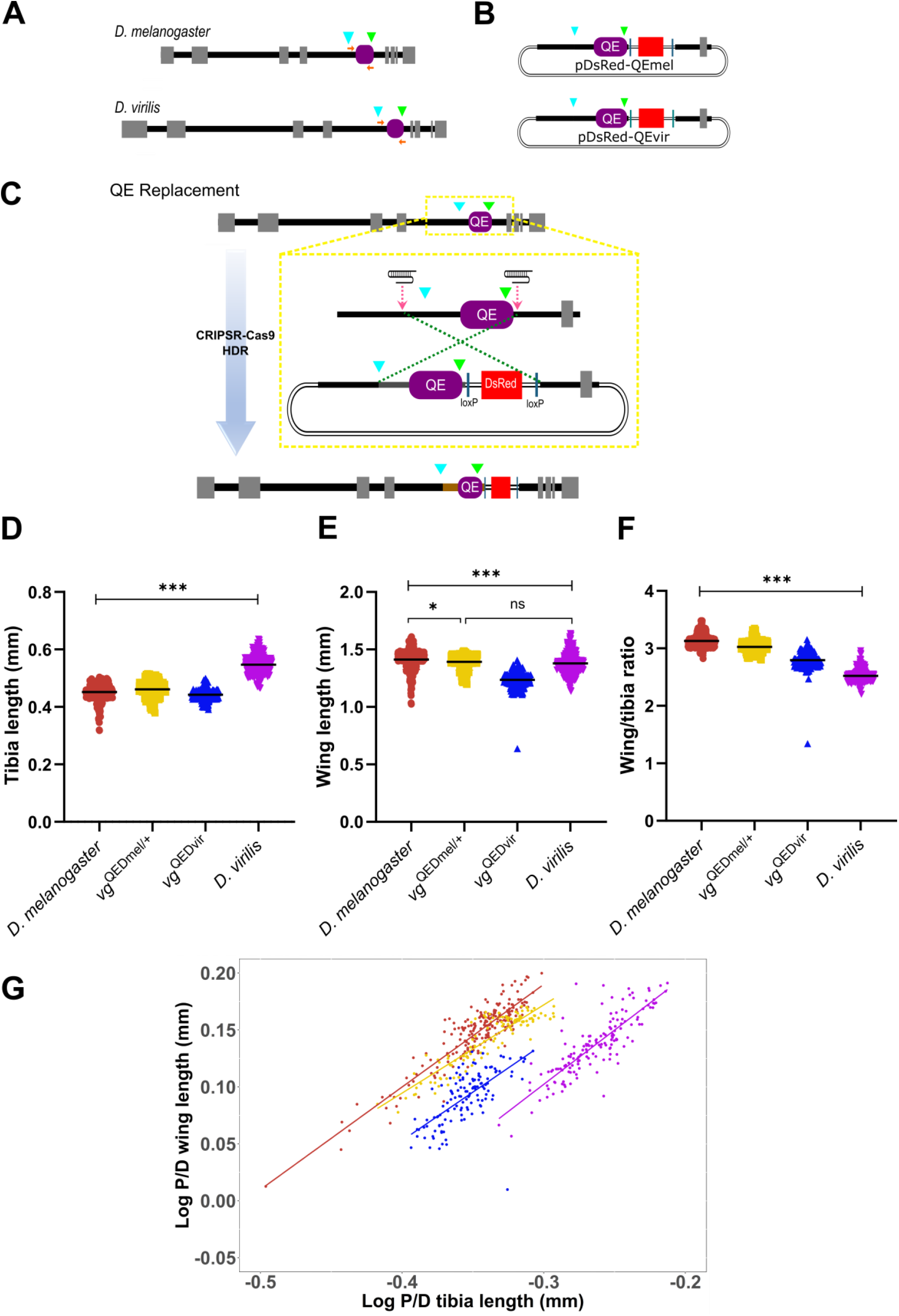
*D. virilis vg*^*QE*^ replacement into *D. melanogaster* results in allometric changes. (A) *vg* gene, QE (purple box) and SID sites (arrowheads) of *D. melanogaster* and *D. virilis* are represented. The amplified and cloned region used for the replacement of the QE in *D. melanogaster* is indicated with the orange arrows. (B) Donor plasmid constructions with QE sequences for each specie. (C) Schematic representation of the *vg*^*QE*^ replacement with CRISPR/cas9 Homologous Directed Repair (HDR) system. (D,E) Proximal-distal lengths of tibias (from anterior legs, D) and wings (E) were measured and compared in female flies (equivalent results were obtained for males; see Fig. S4) in mutant (*vg*^*QEDmel*^, *vg*^*QEDvir*^) and wild type (*D. melanogaster, D*.*virilis)* animals. Each dot in the distribution is the average of each pair of tibias and wings, respectively. (F) Wing-to-tibia ratios from measurements in each side (left and right) of an animal were computed and statistically compared. (D-F) Mean values are shown. One-way ANOVA analysis using multiple comparisons with Tukey test, \*\*\**p<0*.*001, *p=0*.*02, ns =* Not Statistically Significant (*p>0*.*05*). (G) Wing-to-tibia allometric relationships. Yellow and blue lines and dots correspond to HDR replacements using *vg*^*QEDmel*^ and *vg*^*QEDvir*^ respectively. Red and purple lines and dots show the allometric relationship in *D. melanogaster* and *D. virilis* wild type animals respectively. Experiment replicated two times in the laboratory. *D. melanogaster n*= 224, *vg*^*QEDmel*^ *n*=120, *vg*^*QEDvir*^ *n*=128, *D. virilis n*=156.

### CRISPR/Cas9-HDR replacement of the *vg*^*QE*^ from *D. virilis* into the *D. melanogaster* genome

To evaluate the influence of the *vg*^*QE*^ in *Drosophila* wing allometry, we use the CRISPR/Cas9 Homologous Directed Repair (HDR) system (Evans, 2017; Port et al., 2014) to replace the *vg*^*QE*^ sequence in *D. melanogaster* with the predicted *vg*^*QE*^ from other species. We decided to replace only with the *vg*^*QE*^ of *D. virilis* because only in this species we obtained a dramatic change in their wing-to-tibia allometries (Fig. 3D; Fig. S2D). Thus, we designed a guide RNA (gRNA) expression vector using pCFD4 backbone (Howard et al., 2021; Port et al., 2014) that contains two gRNA sequences to target the *vg*^*QE*^ region in *D. melanogaster* (Fig. 5A-C). We then generated a *vg*^*QE*^ homologous donor plasmid that contains the sequence of the predicted *vg*^*QE*^ of *D. virilis* and *D. melanogaster* (as a control) to act as a template for HDR repair system (Fig. 5B). The experimental design of this donor construct allowed us to generate a base vector, that contains left and right *D. melanogaster* homologous arms and DsRed, that could work as a single vector to produce different *vg*^*QE*^ replacements using the same gRNA sequences and donor backbone of pHD-DsRed (Fig. 5C). Plasmids with the gRNAs and the *D. virilis* or *D. melanogaster vg*^*QE*^ vectors were verified through PCR (Figure S3A, B) and injected into flies that maternally carry the Cas9 endonuclease (nos-Cas9) to generate the *vg*^*QE*^ replaced flies.

### Replacement of the *vg*^*QE*^ from *D. virilis* into the *D. melanogaster* genome change the wing-to-tibia allometry

After obtaining stable stocks in which the *vg*^*QE*^ of *D. melanogaster* (*vg*^*QEDmel*^) or *D. virilis* (*vg*^*QEDvir*^) was replaced with the endogenous *vg*^*QE*^ of *D. melanogaster*, we asked if these replacements could affect the wing-to-tibia scaling relationships. With this aim, we generated homozygous stocks (except in the case of *vg*^*QEDmel*^, for which homozygous animals were not viable and we used *vg*^*QEDmel*^/+ flies instead) and plotted allometric relationships as we did for wild-type animals (Fig. 5D-G; Fig. S4A-D). Tibia measurements are only slightly (although significantly) different in *vg*^*QEDmel*^ and *vg*^*QEDvir*^ flies (Fig. 5D; Fig. S4A), but *vg*^*QEDvir*^ animals have a marked reduction in wing length with respect to *vg*^*QEDmel*^ controls, and wild-type *D. virilis* itself (Fig. 5E; Fig. S4B). This suggests that the *vg*^*QE*^ of *D. virilis* likely carries genetic information to underscale wing size. This underscaling effect, however, was not as dramatic as in wild-type *D. virilis*, as revealed by the wing-to-tibia ratios and allometries (Fig. 5F, G; Fig. S4C, D), suggesting that other inputs were needed to achieve the degree of underscaling that *D. virilis* exhibits. Nonetheless, this experiment reveals that a single *cis*-regulatory element has the potential to modulate scaling relationships in animals.

## DISCUSSION

The evolution of scaling relationships between body parts is a major source of phenotypic diversity. The diversity of wing to whole-body relationships among insect species displays an astonishing repertoire of allometries; butterflies have enormous wings compared to their bodies, while bees have quite the opposite wing-to-body ratio. While the evolution of these scaling relationships is likely to be very complex among distant species, the evolution of wing to whole-body scaling in more closely related species may be pinpointed to specific regulatory changes in the genome. The *cis*-regulatory hypothesis, in which phenotypes evolve through changes in *cis*-regulatory regions of key genes has received a lot of attention in the literature (Carroll et al., 2005; Jiggins et al., 2017; Rebeiz and Tsiantis, 2017; Wray, 2003; Wray, 2007); however, very few examples in which evolutionary changes were mapped to specific *cis*-regulatory interactions have been reported (Stern and Orgogozo, 2009; Wittkopp and Kalay, 2011).

In this work, we used CRISPR-mediated replacement (Fig. 5A-C) of a specific regulatory sequence within the *vg* wing selector gene from *D. virilis*, that diverged from *D. melanogaster* about 60 million years ago (Powell and DeSalle, 1995) and displays underscaling of wings relative to the rest of the body (Fig. 3), into the genome of *D. melanogaster*. We found that this *cis*-regulatory itself does contribute, at least partially, to this allometric change (Fig. 5F,G), providing support to the hypothesis that evolution of scaling relationships could be driven by genetic variations in *cis*-regulatory elements (Fig. 1C).

How could the replacement of a *cis*-regulatory element affect the ability to scale an organ with respect to the size of another reference within the animal? The fact that the *vg*^*QEDvir*^ somehow ‘senses’ the size of the animal so that it produces underscaled wings both in the genomes of *D. virilis* and *D. melanogaster* suggests that the Dlp8-mediated inter-organ regulation that has been previously reported in *D. melanogaster* (Boulan et al., 2019; Colombani et al., 2012; Mesquita et al., 2010) could be modulated by this *cis*-regulatory element. In particular, it is possible that the *vg*^*QEDvir*^ contains repressor domains that prevent a wing *vs*. whole-system coordination of growth. This *vg*^*QEDvir*^ repressor hypothesis could also explain why when we delete the *vg*^*QE*^ sequence in *D. melanogaster*, the wing phenotype is fully rescued (Farfán-Pira et al., 2022), presumably through shadow enhancers (Hong et al., 2008), but in the *vg*^*QEDvir*^ replacement is not.

The *vg*^*QE*^ sequence has been studied to determine which components are necessary to induce *vg* transcription, where SID-TEA domains of Scalloped (Sd) interact with DNA consensus sequence (TGGAATGT) located in the QE region (Halder et al., 1998; Klein and Arias, 1999; Simmonds et al., 1998). As we noted in our alignment (Fig. 4), the sequences of the *D. melanogaster, D. simulans, D. ananassae*, and *D. virilis*, SID-TEA domains are evolutionarily conserved, suggesting that this role of Sd to form a complex with Vg is a conserved mechanism to induce the transcriptional mechanism of *vg* expression in *Drosophila*. But, additional sites within the QE also influence *vg* transcription; for example, the *Drosophila* transcription factor Mothers Against Dpp (MAD) has an N-terminal homology region (mad1) that binds within the QE (GCCGnCGC) sequence in *D. melanogaster* and has a role in direct expression of *vg* in the pouch and wing formation. Furthermore, mutation of this consensus sequence and the inhibition of interaction between MAD and the QE sequence, produces animals with smaller wings (Certel et al., 2000; Kim et al., 1997). According to other reports, in *D. virilis* the MAD binding site is present in the sequence of regulatory sequences of the *daughters against dpp* (*dad*) gene (Weiss et al., 2010), suggesting that *D. virilis* may have similar mechanisms of *vg* regulation by MAD in QE sequences, but further studies are necessary to confirm or refute this hypothesis.

Another aspect about *vg* regulation in *D. melanogaster*, is the participation of Wingless (Wg) and Dpp on QE activation (Parker and Struhl, 2020; Zecca and Struhl, 2007b). Previous studies in *D. melanogaster* and *D. virilis* suggest that Dpp has a positive role and Wg has a negative role in the regulation of an enhancer to promote the visceral mesoderm induction (Lee and Frasch, 2005). In addition to these components, a DFR (*Drosophila* Drifter) POU Domain transcription factor has been described in *D. virilis* and *D. melanogaster* that binds to an adjacent site in QE MAD-binding site that is essential to QE activation in the wing pouch, through a complex between MAD and DFR (Certel et al., 2000). This MAD site is apparently also a binding site for Brinker (Brk), presumably acting as a competing repressor of *vg* transcription (Kirkpatrick et al., 2001). Perhaps in the *vg*^*QEDvir*^ replacement, Brk-mediated repression is somehow favored supporting our *vg*^*QEDvir*^ repressor hypothesis described above.

Strikingly, the presence of *vg*^*QE*^ sequences located in the fourth intron of the *vg* gene in the honey bee, *Apis mellifera* (which diverged from *Drosophila* about 300 million years ago) shows similar patterns of expression in imaginal wing discs (Prasad et al., 2016). Thus, the evolutionary conservation of QE in other insect groups suggest that our finding that *cis*-regulatory role of the *vg*^*QE*^ in driving wing scaling relationships may be relevant in more distant species as well.

## Acknowledgements

We thank The Company of Biologists for a Travelling Fellowship grant DEVTF2105555 (https://www.biologists.com/travelling-fellowships/) awarded to Keity J. Farfán-Pira; to José Luis Fernández-López and Rafael Rodríguez-Muñoz for technical assistance; to Consejo Nacional de Ciencia y Tecnología (CONACyT) for PhD fellowship (487729) awarded to Keity J. Farfán-Pira. We thank Eduardo Ochoa for taking and sharing the photos of the specimens in Fig. 1. We also thank Ehab Abouheif, Fanis Missirlis, Augusto Cesar Poot-Hernández and members of the Nahmad and Evans laboratories for discussions.

## Author contributions

Conceptualization: M.N., T.A.E.; Methodology: M.N., T.A.E., K.J.F.P., T.I.M.C.; Formal analysis: M.N., T.A.E., K.J.F.P., T.I.M.C.; Resources: T.A.E.; Data curation: K.J.F.P., T.I.M.C.; Writing-original draft: M.N., K.J.F.P.; Writing-review and editing: M.N., T.A.E., K.J.F.P., T.I.M.C.; Visualization: M.N., K.J.F.P., T.I.M.C.; Supervision: M.N., T.A.E.; Funding acquisition: M.N., T.A.E.

## Data availability

Data are available from GitHub (https://github.com/KeiFarfan/evo-devoDrosophila.git).

